# Self-assembly is important for the target membrane recruitment of a nuclear dynamin-related protein

**DOI:** 10.1101/2025.02.19.639038

**Authors:** Sakti Ranjan Rout, Faiyaz Alam, Swapnil Sahoo, Abdur Rahaman

## Abstract

Dynamin superfamily proteins are large GTPases that perform their cellular functions by self-assembling on their target membranes. Dynamin-related protein 6 (Drp6) associates with the nuclear membrane and performs nuclear remodeling. However, the mechanism of its recruitment to the target membrane is not known. Here, we discover that self-assembly of Drp6 is essential for its nuclear membrane recruitment. We identified four residues, 411-GKFR-414 to be essential for its self-assembly. We also demonstrated that the mutant Drp6 (Drp6_GKFR-AAAA_) failed to recruit to the nuclear membrane. This loss of nuclear membrane recruitment is not due to the lack of membrane binding capability, since the mutated protein was able to bind membrane prepared *in vitro*. Together, our results suggest that in addition to membrane binding, self-assembly of a nuclear dynamin-related protein is also important for the target membrane recruitment.

## Introduction

The dynamins and dynamin-related proteins (DRPs) are a group of large GTPases that perform membrane remodeling functions in various cellular organelles. These multi-domain proteins execute their functions mainly by fission, fusion, or tubulation of the target membrane (Praefcke & McMahon, 2004). The classical endocytic dynamins comprise an N-terminal GTP hydrolyzing domain (GD), a middle domain (MD) involved in oligomerization/self-assembly, a membrane-binding PH domain (PHD), a GTPase effector domain (GED), and a proline-rich domain (PRD) that interacts with accessory proteins (Praefcke & McMahon, 2004). DRPs, a sub-set of the dynamin superfamily, lack PHD and PRD(Chappie & Dyda, 2013; Hong et al., 2003). However, DRPs harbor a domain responsible for membrane binding in place of PHD (Ramachandran & Schmid, 2018). Members of the dynamin superfamily self-assemble on their target membranes, which is a prerequisite for their functions (Hinshaw, 2000). Endocytic dynamins form homodimers in solution, which subsequently interact to form stable tetramers during the initiation of self-assembly. These tetramers translocate to the template membrane and act as a nucleating center for the higher-order multimeric organization (Accola et al., 2002; Carr & Hinshaw, 1997; Sweitzer & Hinshaw, 1998). Although the state of oligomerization varies depending on the members, self-assembly of all known dynamin superfamily proteins requires the participation of GD, MD and GED (Accola et al., 2002; Carr & Hinshaw, 1997; Chang et al., 2010; Daumke et al., 2010; Hu & Rapoport, 2016; Nigg & Pavlovic, 2015; Ramachandran et al., 2007; Sever et al., 2006). One of the accepted models suggests that the C-terminal-GED helix of one dynamin polypeptide interacts in trans with the G domain of another polypeptide, forming a domain-swapped dimer which subsequently associates through their stalk regions to generate a tetramer (Chappie et al., 2011; Pawlowski, 2010). These tetramers in the cytosol are assembly-incompetent but can bind to target membranes via the membrane-binding domain, upon which they undergo a conformational shift exposing the higher order assembly interface, thereby triggering the rapid and cooperative formation of a helical dynamin collar at target sites (Chappie et al., 2011; Fribourgh et al., 2014; Gao et al., 2010). The members of this family share three distinct levels of assembly interfaces that include dimeric interface (interface 2), the oligomeric interfaces (interfaces 1 and 3), and the higher-order helical assembly interface (interface 4) (Alvarez et al., 2025).

The dynamin-related protein 6 (Drp6) associates with the nuclear envelope and regulates nuclear remodeling in *Tetrahymena thermophila* (Rahaman et al., 2008). Recently, it has been demonstrated that a lipid binding domain is required for membrane association and nuclear recruitment of Drp6 via cardiolipin interaction (Kar et al., 2021). An earlier study had shown that Drp6 self-assembles into higher-order oligomeric structures independent of a template membrane (Kar et al., 2018a). However, the role of self-assembly in nuclear membrane recruitment of Drp6 is not known. In this study, we have identified an interface important for self-assembly and evaluated the role of self-assembly in the nuclear recruitment of Drp6. Our results suggest that self-assembly, though not essential for cardiolipin-mediated membrane binding, is important for nuclear membrane association.

## RESULTS

### Identification of a Self-assembly interface in the nuclear dynamin-related protein Drp6

Drp6, like other members of the dynamin superfamily, undergoes oligomerization and self-assembly (Kar et al., 2018a). Sequence alignment of DRP6 with various family members identified a potential self-assembly interface comprising a stretch of four amino acid residues (411-GKFR-414) which aligned with the known interface of other members including MxA (YRGR), MxB (YRGK), hDyn1(IHGIR) and Drp1 (GPRP) (Figure 1a) (Faelber et al., 2011; Fribourgh et al., 2014; Fröhlich et al., 2013; Gao et al., 2010). Based on this result, we have mutated these four residues to alanine (Drp6_GKFR-AAAA_) and named it Drp6-4A. Drp6-4A and the wildtype Drp6 (Drp6) were separately expressed as N-terminal histidine-tagged proteins in *E. coli*. The expressed proteins were purified using Ni-NTA affinity purification, and the purity was checked using Coomassie-stained SDS-PAGE analysis. Drp6 was estimated to be ∼80% pure, and Drp6-4A to be ∼65% pure (Figure 1b). The identity of the purified proteins was confirmed by western blot analysis using anti-His antibody (Figure 1b). Both Drp6 and Drp6-4A were subjected to size exclusion chromatography. Drp6 eluted primarily (∼60%) in the void volume (peak1, 7.8 ml) as higher-order structures (Figure 2). Considering the exclusion limit of the column 660 kDa, these structures comprise at least six protomers. The remaining protein eluted at around 10.3 ml (peak2, ∼ 10%) and 13.2 ml (peak3, ∼30%), corresponding to the tetramers and monomers of Drp6, respectively (Figure 2). This result confirms our earlier observation that Drp6 self-assembles to form higher-order structures in solution. Interestingly, we observed a three fold reduction in the population of self-assembled Drp6-4A (Peak1) as compared to that of Drp6, with a concomitant two fold increase in tetrameric (peak2, ∼19%) and monomeric (peak3, ∼58%) peaks (Figure 2). The presence of the protein in the corresponding peaks was confirmed by western blot analysis using anti-His antibody (Figure 2). We did not consider peak4 (eluted after 16 ml) since it did not contain protein of interest, as confirmed by western blot analysis (Figure 2). Moreover, the calculated molecular weight (∼30 kDa) of this peak is smaller than that of a Drp6 monomer (∼87kDa). These results suggest that the amino acid residues GKFR form an interface important for the self-assembly of Drp6. To further confirm the disruption of Drp6-4A self-assembly, we performed transmission electron microscopy (TEM) after negative staining and compared it to that of Drp6. As reported earlier, Drp6 readily formed self-assembled structures appearing as rings and helical spirals (Figure 3). However, analysis of more than 200 electron micrographs of Drp6-4A failed to identify the presence of any such self-assembled structures (rings and helical spirals), instead, small particles were observed in all the micrographs (Figure 3). Therefore, it can be concluded that the four amino acid residues (411-GKFR-414) are important for Drp6 self-assembly.

**Figure 1:**
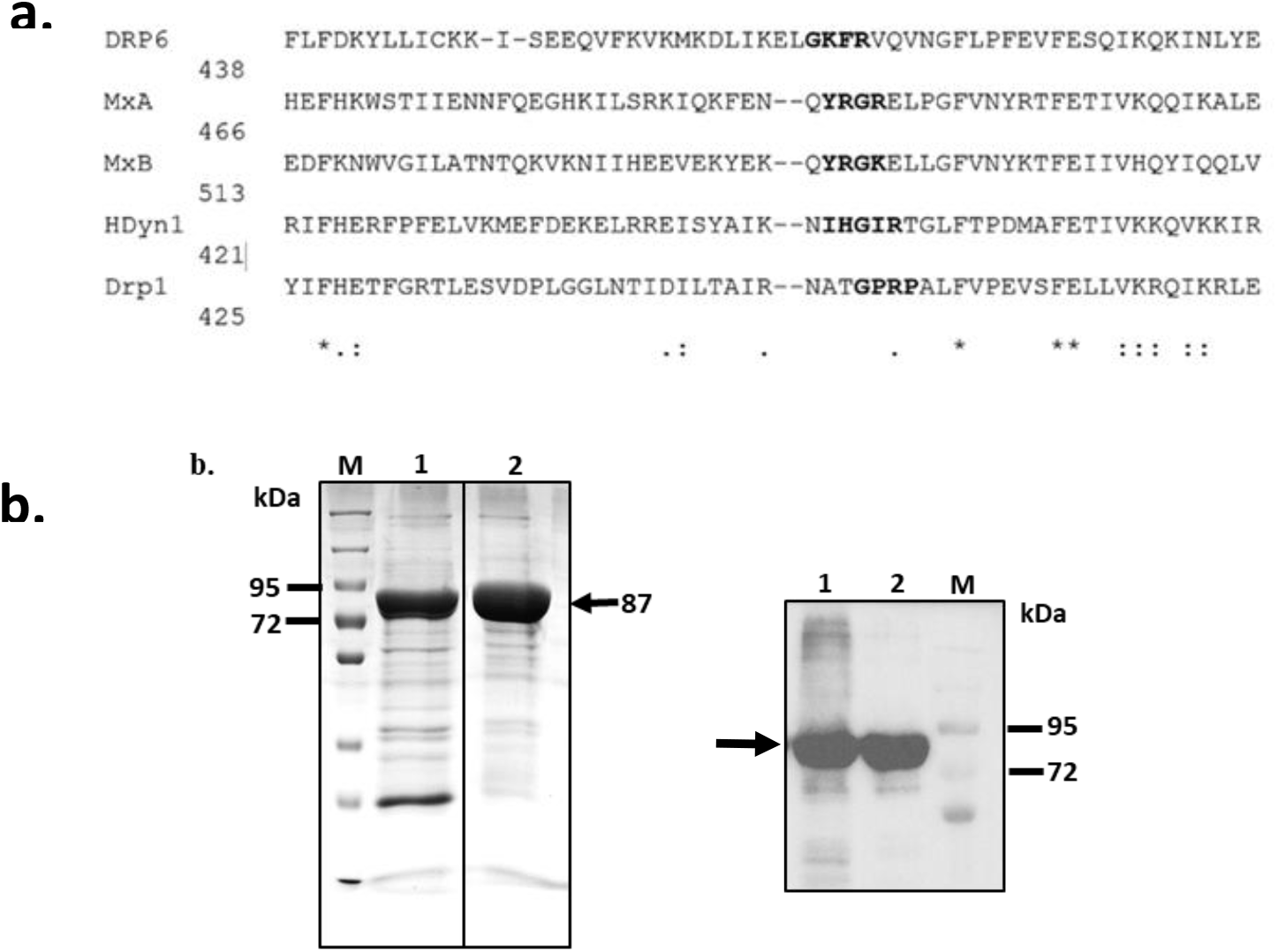
Expression and purification of a putative self-assembly interface mutant. a. Amino acid sequence alignment of Drp6 with other dynamin family members using Clustal Omega (https://www.ebi.ac.uk/jdispatcher/msa/clustalo, Goujon et al., 2010; Sievers et al., 2011). The interface shown to be important for self-assembly in hdyn1, Drp1, MxA and MxB is highlighted in bold letters. A stretch of four amino acids, GKFR is identified as potential self-assembly interface in Drp6. b. Left panel: Coomassie stained SDS-PAGE gel showing purified His-Drp6-4A (lane 1) and His-Drp6 (lane 2). M denotes molecular weight markers, two of which are indicated by a line on the left. The position of the proteins are also indicated by an arrow on the right. Right panel: Western blot analysis of His-Drp6-4A (lane1) and His-Drp6 (lane 2) using anti-His monoclonal antibody (1:5000). The position of the purified proteins are indicated by an arrow.

**Figure 2:**
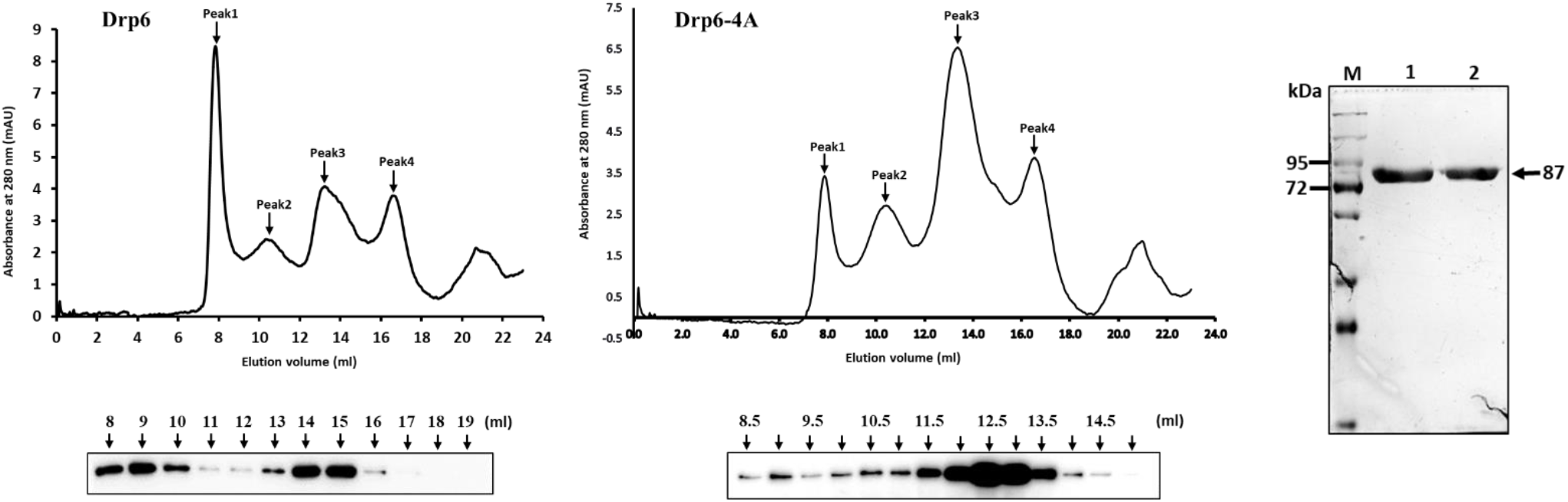
Mutation of the putative self-assembly interface disrupts higher order oligomerization of Drp6. Chromatograms depicting elution profiles of His-Drp6 (left) and His-Drp6-4A (middle) using Superdex-200 size exclusion column. The various peaks are indicated in which elution volume of peak 1 coincides with the void volume. Peak 2 and peak 3 corresponds to tetramer and monomer respectively as estimated from the molecular weight markers. The presence of the protein in the alternate fraction (∼500 µl each) of Drp6 and in all the fractions of Drp6-4A is checked by western blot analysis and shown at the bottom of the chromatograms. Silver stained SDS-PAGE gel (right) of the pooled fractions from peak 1 to peak 3 of both Drp6 and Drp6-4A. Presence of only the specific band confirms that these peaks arise from the protein of interest, and not due to other contaminating proteins.

**Figure 3:**
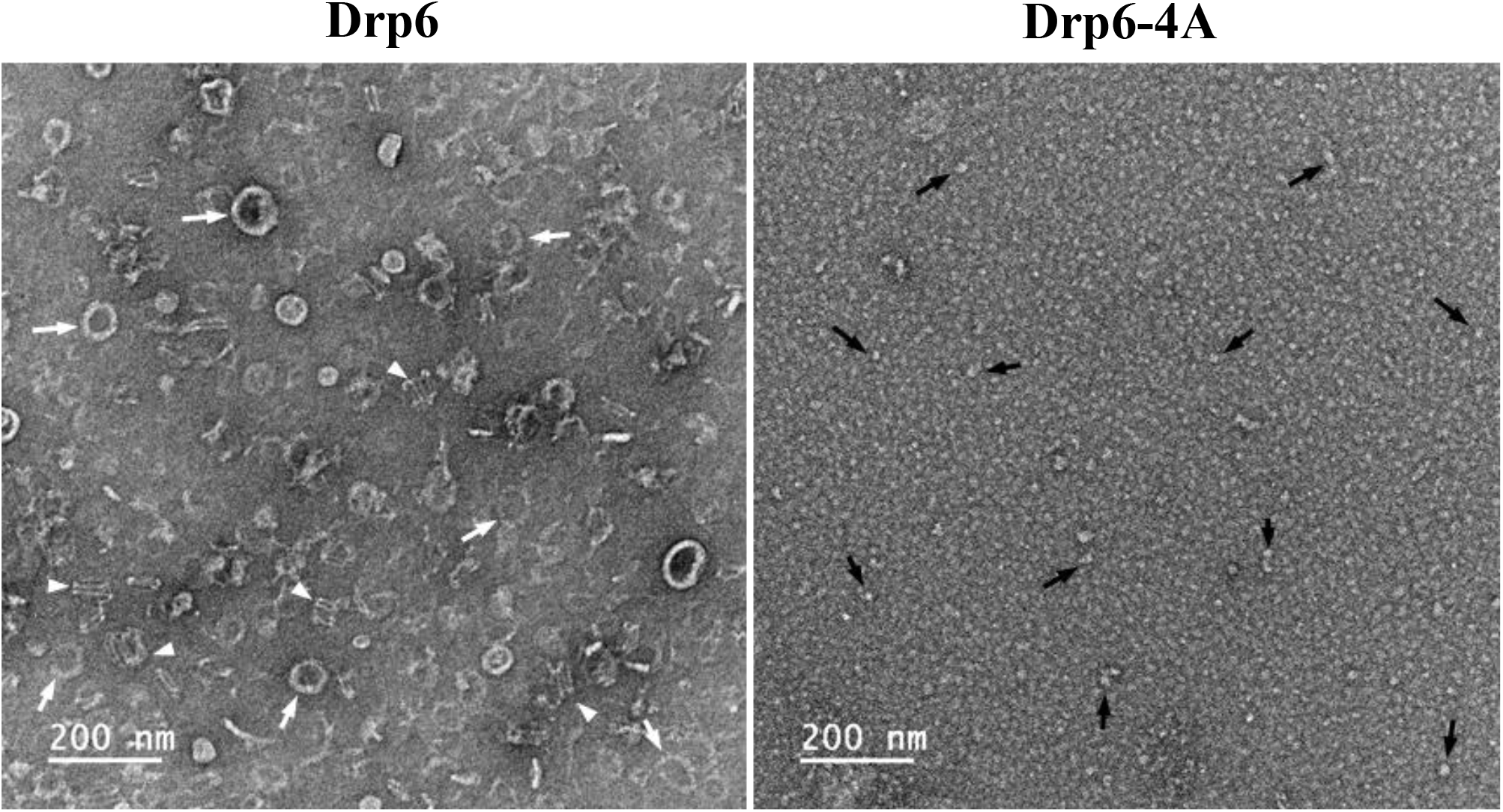
Transmission electron microscopy confirms loss of self-assembled structures in the Drp6-4A. Electron micrographs of negatively stained His-Drp6 (left) and His-Drp6-4A (right) at two different magnifications. The pooled fractions from the gel filtration columns were used for the experiment. Helical spirals and the ring structures are found only in His-Drp6 and are indicated by arrow head and white arrow respectively. Some of the small particles in Drp6-4A are also indicated by arrow.

### Self-assembly of Drp6 is required for GTPase activity and nuclear envelope recruitment

Stimulation of GTPase activity of all known dynamins depends on their self-assembly (Warnock et al., 1996). Since DRP6-4A shows disrupted self-assembly, we hypothesized that this mutation may result in defective GTPase activity. In an *in vitro* GTPase activity assay while DRP6 hydrolyzed GTP at a rate of 2.57 ± 0.56 μM/μM/min, Drp6-4A showed an approximately six-fold reduction in its GTPase activity (0.83 ± 0.01 μM /μM/min) (Figure 4a). This result suggests that, similar to other family members, the stimulation of GTPase activity of Drp6 is dependent on its self-assembly.

**Figure 4:**
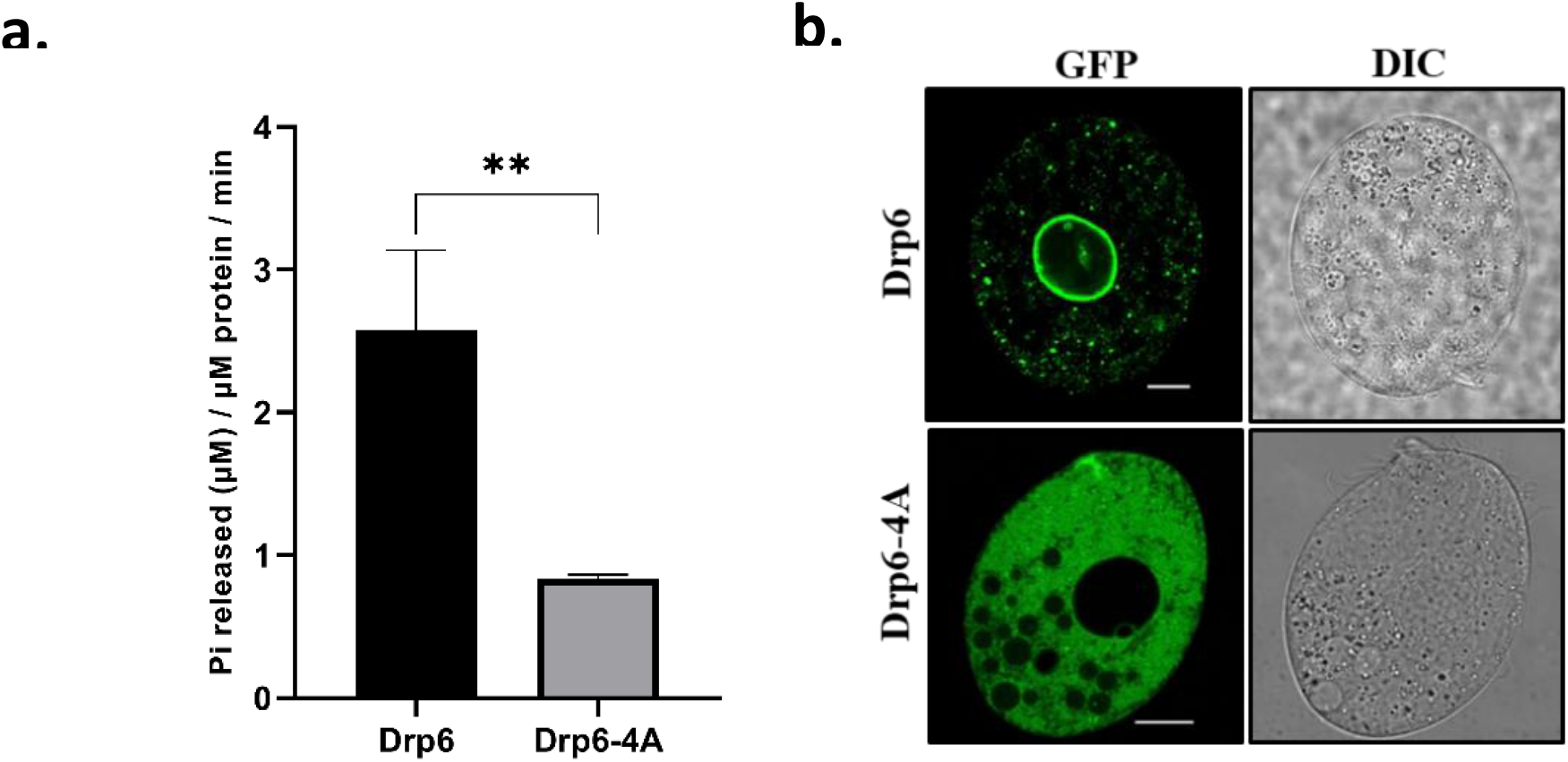
Mutation in the self-assembly interface inhibits GTPase activity and abrogates nuclear localization of Drp6. a. GTP hydrolysis activity of His-Drp6 and His-Drp6-4A. The reaction was carried out for 20 min and the amount of Pi released was estimated and plotted. The statistical analysis was performed using unpaired t-test and the difference was found to be significant (p=0.0061), n = 9. b. Confocal images of live *Tetrahymena* cells expressing either GFP-Drp6 (top) or GFP-Drp6-4A (bottom). Corresponding DIC images are shown on the right. Bar = 10 µm

The dynamin family proteins perform membrane remodeling functions by associating with their target membranes. Drp6 is earlier shown to undergo continuous association and dissociation on the nuclear membrane (Rahaman et al., 2008). To examine if Drp6 association with nuclear membrane prerequisites self-assembly, we expressed Drp6 and Drp6-4A separately as N-terminal GFP fusion proteins and analyzed their localization by confocal microscopy. As expected, Drp6 is predominately present on the nuclear envelope (Figure 4b), confirming our earlier observation. However, we did not detect any enrichment of the mutant Drp6-4A on the nuclear membrane (Figure 4b). This result clearly suggests that Drp6 self-assembly is important for its recruitment to the target membrane.

### Loss of nuclear localization of self-assembly defective Drp6 mutant (Drp6-4A) is not due to loss of cardiolipin interaction

As reported earlier, the recruitment of Drp6 to the nuclear membrane depends on its interaction with cardiolipin (Kar et al., 2021). To test if the loss of nuclear recruitment of Drp6-4A is also due to the loss of its interaction with cardiolipin, we performed an *in vitro* membrane binding assay using recombinant proteins and cardiolipin (CL) containing liposomes. As shown in the figure 5, Drp6 floated to the top fractions, confirming its interaction with Cardiolipin. Interestingly, when Drp6-4A mutant protein was used in a similar assay, it also floated to the top fractions (Figure 5) suggesting that the mutation in the self-assembly interface does not result in the loss of interaction with CL. Therefore, it can be concluded that the loss of nuclear recruitment of Drp6-4A is not due to the loss of cardiolipin interaction. Moreover, Drp6 is also known to interact with two other lipids, namely phosphatidyl serine (PS) and phosphatidic acid (PA) (Kar et al., 2021). To evaluate if the mutant retains its interaction with these two lipids, we performed a similar assay using either PS or PA liposomes. Similar to Drp6, Drp6-4A also co-floated to the top fractions when incubated with these liposomes (Figure 5). These results show that self-assembly is not essential for binding to membranes containing any of these lipids. Therefore, the loss of Drp6 recruitment to the nuclear envelope is not due to its inability to bind the membrane. Taking these results together, we conclude that in addition to cardiolipin binding, nuclear recruitment of Drp6 also depends on its self-assembly.

**Figure 5:**
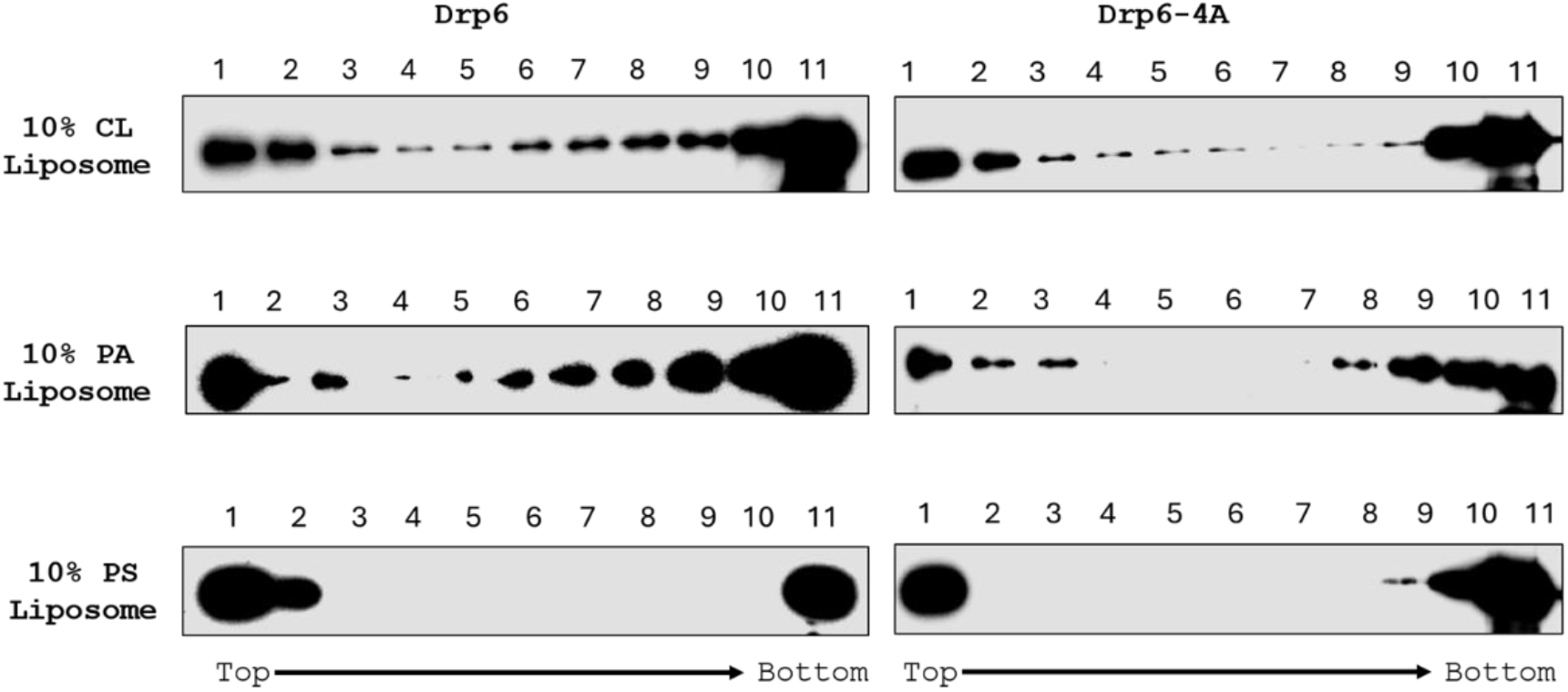
Drp6-4A retains its interaction with the membranes containing either CL, PA and PS. Western blots showing different fractions (top to bottom) of floatation assays with liposomes containing 10% CL or 10% PA or 10% PS using His-Drp6 (left panels) and His-Drp6-4A (right panels).

## Discussion

Members of the dynamin superfamily proteins self-assemble on their target to perform membrane remodeling (Chappie & Dyda, 2013; Fröhlich et al., 2013b; Song et al., 2004). Drp6 regulates nuclear remodeling in *Tetrahymena* and undergoes cycles of assembly/disassembly on the nuclear membrane (Kar et al., 2018a, 2021). In the present study, we have identified the self-assembly interface and assessed the role of self-assembly in the nuclear recruitment of Drp6. The results show that the self-assembly of Drp6 is important for its nuclear membrane association.

Self-assembly is the hallmark of all dynamin proteins. Detailed structural studies in several dynamin members have shown the presence of similar interfaces required for oligomerization and higher-order self-assembly (Chappie & Dyda, 2013; Ford et al., 2011; Gao et al., 2010b; Reubold et al., 2015; von der Malsburg et al., 2011). Importantly, mutations in interface 3 disrupt the self-assembly of many members including endocytic dynamins, MxA, and MxB (Fribourgh et al., 2014; Gao et al., 2010). The presence of the same self-assembly interface, interface 3 in Drp6, reiterates the similarity in its overall structure.

Members of the dynamin family show high basal GTPase activity, which stimulates many folds upon self-assembly on the target membrane (Warnock et al., 1996). Although Drp6 showed stimulation of GTPase activity upon self-assembly, the basal GTPase activity was negligible, and the stimulated activity (2.41 μM/μM/min) was only comparable to the basal activity of dynamins (1.75 μM/min; Song et al., 2004). These results indicate a different mechanism of function for Drp6 since high GTPase activity is important for the functions of other members. Similar to Drp6, Mfn2 also exhibits slow GTPase activity. Recent structural studies have shown that the slow GTPase activity of Mfn2 is due to the sustained stability of the G domain interface (Li et al., 2019). It is possible that the slow GTPase activity of DRP6 may also result from a similar structural arrangement. However, a detailed structural study is required to establish the molecular basis for the slow GTPase activity of DRP6.

Dynamin family proteins are generally associated with the target membrane as either dimers or tetramers and self-assemble upon membrane binding (Jimah & Hinshaw, 2019). However, the self-assembly of Drp6 does not depend on membrane binding (Kar et al., 2018). This result suggests that members of dynamin family proteins show differences in self-assembly requirements for target membrane recruitment. Although majority of the dynamin family proteins do not depend on self-assembly for target membrane recruitment, Drp1 self-assembles in solution and is recruited to the target membrane in its self-assembled form (Chang et al., 2010). Similarly, Drp6 also readily self-assembles in solution, even in the absence of a membrane. The present study shows that the loss of self-assembly results in the loss of Drp6 recruitment to the nuclear membrane. Therefore, like Drp1, the self-assembly of Drp6 is important for binding to the target membrane. Moreover, another dynamin MxA forms a self-assembled structure without binding to its target. Though this self-assembled pool of MxA is proposed to be a storage form, which upon activation signal disassembles for its target binding, there is no experimental evidence to support this proposition (Accola et al., 2002; Daumke et al., 2010; Nigg & Pavlovic, 2015). Hence, we propose that a subset of dynamin-related proteins, including Drp6, Drp1, and MxA, depends on self-assembly for their target recruitment.

As reported earlier, Drp6 binds to the target membrane via interaction with cardiolipin, and inhibition of this interaction results in the complete loss of its nuclear localization (Kar et al., 2021). Surprisingly, the loss of nuclear association of self-assembly defective mutant does not abrogate cardiolipin binding. Therefore, the loss of nuclear membrane recruitment of the self-assembly defective mutant of Drp6 is not due to its loss of interaction with cardiolipin. All the results discussed here suggest that membrane binding is not sufficient, but self-assembly of a nuclear dynamin-related protein is also essential for its recruitment to the target membrane.

## MATERIALS AND METHODS

### Cloning, expression, and purification of Drp6 and Drp6-4A

Drp6 in pRSETb, previously cloned at KpnI and EcoRI sites (Kar et al., 2018b) was used as a template to generate Drp6-4A (Drp6_GKFR-AAAA_) with 5’-ATGAAAGATCTGATTAAAGAACTGGCCGCAGCTGCAGTGCAGGTGAACGGCTTT CTGCCG-3’ as forward primer and 5’-CGGCAGAAAGCCGTTCACCTGCACTGCAGCTGCGGCCAGTTCTTTAATCAGATCT TTCAT-3’ as reverse primer following Quick Change Protocol (Stratagene). Drp6 and Drp6-4A in pRSETb were transformed into C41 (DE3) *E. coli* strain and expressed as N-terminal His_6_ tag proteins. The expression was performed with either 0.5 mM or 0.25 mM IPTG for Drp6 or Drp6-4A, respectively, at 18°C for 16 h. The cells were harvested at 19,000 g for 10 min at 4°C, and the pellet was re-suspended in lysis buffer (25 mM HEPES pH 7.5, 300 mM NaCl, 2 mM MgCl_2_, and 2 mM β-ME) supplemented with 10% glycerol, 2 μg/ml DNase I (Sigma), 200 μg/ml lysozyme and EDTA free protease inhibitor cocktail (Roche). The cells were lysed 1 h post-incubation by sonication, and the cell lysate was clarified by centrifugation at 24,652 g for 45 min at 4°C. The clarified lysate was incubated with Ni-NTA resin for 1 h at 4°C and the resin was washed with 100 bed-volume of ice-cold lysis buffer supplemented with either 50 mM imidazole for DRP6 or with 40 mM imidazole for DRP6-4A. The bound protein was eluted with 250 mM imidazole in lysis buffer. The purified protein was analyzed by SDS-PAGE and western blot using anti-His monoclonal antibody (1:5000, Sigma).

For expression in *Tetrahymena thermophila*, the Dpr6-4A was PCR amplified using Drp6-4A in pRSETB as template, 5’-CACCCTCGAGATGACCAACACCATTGTG-3’ as forward primer, and 5’-GCGGGGCCGAATTCTCAATCA-3’ as reverse primer and the amplified product was introduced into pIGF vector using Gateway cloning strategy (Invitrogen) to express it as N-terminal-GFP fusion protein. Conjugating wild-type *Tetrahymena* cells were transformed with the constructs by electroporation, and the transformants were selected using 100 µg/ml paromomycin sulfate (PMS) in SPP media. Cells were grown to a density of 1 × 10^5^ cells/ml, and expression was induced by adding cadmium chloride (1 µg/ml) for 3 h before analyzing by confocal microscopy using Leica DMI8 confocal microscope.

### Size Exclusion Chromatography

Size exclusion chromatography was performed in Superdex 200 GL 10/300 column (GE Life Sciences, USA) using Akta Explorer FPLC system (GE Healthcare, USA). The column was washed with 2 column-volume (CV) of autoclaved Milli-Q water before equilibration with buffer B (25 mM HEPES pH 7.5, 150 mM NaCl, and 2 mM MgCl_2_). Five hundred microliters of protein (0.5 mg/ml) was injected into the pre-equilibrated column and was run at 0.5 ml/min flow rate. The chromatogram was recorded by taking absorbance at 280 nm. Fractions of 0.5 ml each were collected and checked by western blot analysis using anti-His monoclonal antibody (Sigma).

### Flotation Assay

Liposomes (2.5 mg/ml) containing 70% PC and 20% PE with either 10% CL, 10% PA, or 10% PS were prepared by dissolving lipids (Avanti Polar) in analytical grade chloroform in a round bottom flask. The solvent was then evaporated in a lyophilizer to obtain a thin, dry film. Liposomes were made by resuspending the film in buffer B at 37°C. The resuspended solution was extruded 23 times through an extruder (Avanti Polar) using a 100 nm filter. The size distribution and homogeneity were measured by dynamic light scattering in Malvern Zetasizer Nano instrument (Malvern Panalytical Ltd, UK). For floatation assay, 1 μM protein was incubated with 0.5 mg liposomes in GTPase assay buffer supplemented with 1 mM GTP (Sigma, USA) at room temperature for 1 h. The reaction mixture was mixed with sucrose (final concentration 40%) and placed at the bottom of a 13.2 ml ultra-centrifugation tube. This was overlaid with 2 ml each of 35%, 30%, 25%, 20%, and 1 ml of 15% and 10% sucrose solutions in the same buffer. The gradient was subjected to ultra-centrifugation at 35,000 RPM for 16 h at 4°C using a SW41 rotor in a Beckman Coulter ultracentrifuge (Beckman Coulter Life Sciences, USA). Fractions (1 ml each) were collected from the top, and the presence of protein was detected by western blot analysis using anti-His monoclonal antibody (Sigma).

### GTPase Assay

GTP hydrolysis activity was assessed by a colorimetric assay using BIOMOL Green (ENZO Lifesciences, USA). The reaction (20 μl) was carried out with 1μM protein using 1 mM GTP (Sigma, USA) in buffer B for 30 min at 37°C. The reaction was stopped by adding 5 μl of 0.5 M EDTA, and absorbance was measured at 620 nm. The experiment was repeated thrice in triplicate, and the results were analyzed using GraphPad Prism7 software. An unpaired t-test was performed for statistical analysis.

### Electron microscopy

For this purpose, the protein was purified by Ni-NTA affinity column followed by size exclusion chromatography. The purified Drp6 or Drp6-4A (1 μM) protein was incubated with 0.5 mM GTPγS (Sigma) in buffer B for 20 min at room temperature and was adsorbed onto a 200 mesh carbon-coated Copper grid (Ted Pella, Inc, USA) for 2 min. The grid was then stained with 2% uranyl acetate (MP Biomedicals, USA) for 2 min and was allowed to dry at room temperature before collecting electron micrographs using JEOL JEM F200 Transmission Electron Microscope (TEM) (JEOL, USA)

## Acknowledgements

We thank Dr Utpal Nath from Indian Institute of Science for theirvaluable comments on our manuscript. We also thank members of the lab, Mr Soham Mukhopadhya, Ms Gargi Dey, Mr Kaustubh Prakash and Ms Urbi Ghosh for other helps. We also thank Center for Inter-disciplinary Science, National Institute of Science Education and Research for TEM facility.

